# Multifactorial motor behavior assessment for real-time evaluation of emerging therapeutics to treat neurologic impairments

**DOI:** 10.1101/668327

**Authors:** Riazul Islam, Carlos Cuellar, Ben Felmlee, Tori Riccelli, Jodi Silvernail, Suelen Lucio Boschen, Peter Grahn, Igor Lavrov

## Abstract

Integrating multiple assessment parameters of motor behavior is critical for understanding neural activity dynamics during motor control in both intact and dysfunctional nervous systems. Here, we described a novel approach (termed Multifactorial Behavioral Assessment (MfBA)) to integrate, in real-time, electrophysiological and biomechanical properties of rodent spinal sensorimotor network activity with behavioral aspects of motor task performance. Specifically, the MfBA simultaneously records limb kinematics, multi-directional forces and electrophysiological metrics, such as high-fidelity chronic intramuscular electromyography synchronized in time to spinal stimulation in order to characterize spinal cord functional motor evoked potentials (fMEPs). Additionally, we designed the MfBA to incorporate a body weight support system to allow bipedal and quadrupedal stepping on a treadmill and in an open field environment to assess function in rodent models of neurologic disorders that impact motor activity This novel approach was validated using, a neurologically intact cohort, a cohort with unilateral Parkinsonian motor deficits due to midbrain lesioning, and a cohort with complete hind limb paralysis due to T8 spinal cord transection. In the SCI cohort, lumbosacral epidural electrical stimulation (EES) was applied, with and without administration of the serotonergic agonist Quipazine, to enable hind limb motor functions following paralysis. The results presented herein demonstrate the MfBA is capable of integrating multiple metrics of motor activity in order to characterize relationships between EES inputs that modulate mono- and polysynaptic outputs from spinal circuitry which in turn, can be used to elucidate underlying electrophysiologic mechanisms of motor behavior by synchronizing these datasets to metrics of movement and behavior. These results also demonstrate that proposed MfBA is an effective tool to integrate biomechanical and electrophysiology metrics, synchronized to therapeutic inputs such as EES or pharmacology, during body weight supported treadmill or open field motor activities, to target a high range of variations in motor behavior as a result of neurological deficit at the different levels of CNS.

## Introduction

Damage to the central nervous system (CNS), due to either acute events such as cerebral ischemia or spinal cord injury (SCI), or prolonged neurodegenerative diseases such as Parkinson Disease (PD) or Multiple Sclerosis (MS), can result in lifelong impairment of sensorimotor functions (e.g., reaching, grasping, standing, and/or walking). Once damages, the CNS undergoes a cascade of complex changes both within the brain and across spinal cord sensorimotor networks that integrate sensory signaling and motor control commands to produce coordinated neuromuscular activity. Neuromodulation has emerged over the last decades as a clinically available therapeutic to alleviate neurologic deficits, such as gait dysfunction due to PD^1–4^, tremor^5^, medically refractory central pain syndromes^6^,, dystonia^7^ and epileptic seizures^8^. Additionally, over the last few years, investigations using EES have shown it hold great potential to improve function in humans with spasticity^9,10^ and restoration of motor control after spinal cord injury^11–14^.

One of the most critical concepts of spinal cord neuromodulation is a central pattern generation (CPG) that describes the organization of distinct neural networks that receive patterned sensory signaling and, in turn, produce predictable patterns of motor activity^15,16^. The anatomical overlap of sensory signals that originate in the peripheral nervous system and converge upon spinal sensorimotor networks via multi-segmental innervations to the spinal cord and the brain suggests that, multiple CPGs across central nervous system could integrate sensory signaling to allow real-time optimization of motor outputs in response to external perturbations^15–18^ (Fig. 1a). It is expected that CPGs at different level of CNS are able to coordinate activity during complex motor functions, such as standing, stepping, or running. However, the distinct role of the neuronal circuitry of CPG during motor task performance, as well as the mechanism by which multiple CPGs coordinate their activity remains undefined^19–22^.

Targeting dorsal spinal cord structures and activation spinal CPG, epidural electrical stimulation (EES) has shown promise to restore functions after paralysis due to SCI^11–14^. Using rodent models of either partial or complete SCI, significant progress has been made in understanding underlying mechanisms by which EES alters spinal sensorimotor activity to enable functions^23–30^. The majority of this work has focused on interpreting electromyography recordings from EES-enabled muscle activity synchronized to biomechanical assessments of movements, however, a gap in knowledge remains with respect to direct modulation of spinal sensorimotor activity in relation to end organ function. This gap is largely due to a lack of affordable *in vivo* tools capable of modulating and recording EES-enabled sensorimotor activity synchronized to each electrical pulse that is delivered to neuronal structures, and synchronized to biomechanical assessments of motor performance, all of which are correlated to sensory input to the CNS during different motor patterns.

Several experimental models have been established to estimate interactions between the electric fields that are produced during stimulation and neural tissues in close proximity to stimulation^31,32^. Although, these studies revealed the anatomical structures most likely to be facilitated by EES, and the roles these structures play in producing both desirable and undesirable outcomes, *in silico* properties of computational modelling and limitations of *in vitro* models fail to describe key observations made during *in vivo* investigations, such as the degree to which different parameters of stimulation influence motor output via activation of distinct sensorimotor networks, the impact of pharmacological dosing on network activity, or which motor training-induced network changes are responsible for improvements in motor task performance over the course of EES. Currently available *in vivo* models of neurologic damage lack approaches to investigate spinal sensorimotor network inputs and outputs simultaneously. Additionally, outcomes generated by available models are typically captured using isolated, nonsynchronized, assessments of motor task performance and electrophysiological metrics^24,26,28,33^. Multiple studies have shown that motor training following SCI reorganizes neural circuitry via repetitive reinforcement of sensorimotor network activation patterns^34,35^. For example, in rodents, step training on a treadmill following SCI induces reorganization of spinal sensorimotor ensembles via selective reinforcement of locomotor networks ^26^. Furthermore, step training combined with EES, and/or pharmacological neuromodulation, both of which alter spinal network excitability, enhances motor performance^23^. Positive outcomes from task-specific training in humans after incomplete SCI led to the development of several body weight support systems (BWSs) to achieve appropriate posture in animal models and humans during treadmill training^36–38^ or over-ground locomotion^39^. These BWSs were developed to maximize the effect of task-specific motor training while assessing general aspects of motor performance, however, these systems lack the capability to integrate data recordings across multiple dynamic variables such as limb loading, kinematics, and timing of exogenous neuromodulatory application (e.g., electrical pulse frequency, intensity, pharmacological dosage, time of application). Additionally, mechanisms responsible for locomotion network plasticity in neuromotor impairments (e.g. SCI, PD, tremor, spasticity, MS), as well as the progression of plasticity over the course of motor training, have yet to be elucidated. To our knowledge, currently available assessments of motor performance in rodents are not capable of simultaneously recording and integrating critical parameters to gain knowledge on the mechanisms through which neuromodulation with motor training enables different types of motor behavior and re-organizes spinal networks.

In order to evaluate impaired neural circuitry facilitated by neurostimulation combined with motor-training, we proposed a multifactorial behavioral assessment (MfBA) and designed a new BWS assessment system, which integrates multiple behavioral and electrophysiological parameters and allows simultaneous recording of sensorimotor network inputs and outputs, synchronized to multiple metrics of motor behavior (Fig. 1 a-b). Proposed MfBA approach and system’s capabilities were tested using neurologically intact rodents, parkinsonian (PD), and SCI rodent models that exhibit distinct motor dysfunctions related to the brain and spinal cord lesion. Following validation of proposed MfBA approach in healthy animals, PD, and SCI rats, *we hypothesized that, during functional tasks in SCI rats, EES and pharmacology enabled modulation of mono- and polysynaptic components of spinal circuitry is reflective of changes in motor outputs and animal behavior.*

## Materials and methods

### Multi-factorial behavioral assessment system configuration

The multi-factorial assessment system consists of the following main components: (1) BWS apparatus integrated with force and torque transducers, (2) motion tracking system, (3) open field camera, (4) interface with chronically implanted electrophysiological recording electrodes, (5) interface with chronically implanted neural stimulation electrodes, (6) motorized treadmill, and (7) open field platform. Through hardware level communication between these modules the system combines behavior, locomotion and electrophysiology in either an open field or treadmill environment with the flexibility to position animals for bipedal or quadrupedal stepping. This system is capable of synchronizing input variables including EES, pharmacology, treadmill speed, variation of load, direction of locomotion, and extent of BWS.

#### Body weight support system (BWS)

The BWS system (Fig. 1b, d) provides trunk support to motor impaired rodents while imposing minimal friction forces in order to provide a neutral environment in which to study motor behavior. Rodents were secured to the BWS using a custom-made, fully adjustable fabric padded jacket, to provide BWS during training and assessment activities on a treadmill or in open field. Components of the BWS system were chosen with the intent to achieve friction forces that allow unrestricted motor behavior in the open field environment while also considering overall system cost. The total cost to build the BWS was ~$400 (Sup. table 1 for cost details). The major components of the BWS are described herein:

#### Aluminum frame structure

The BWS structure consists of rectangular aluminum alloy extrusion bars and corner plates (47065T503, Mcmaster-Carr, Elmhurst, Illinois) (Fig. 1b-c). The rigid framework is easily modifiable to accommodate assessment equipment required during motor training and data recording, such as camera angle and height adjustment to during treadmill or open field motion analysis. The goal of the frame is to provide consistent placement of the equipment and be adaptable for new experiments. To ensure that open field test can be performed in the system, the frame was constructed to provide a 120cmx120cmx60cm unobstructed environment.

#### Two axis linear motion

To allow for planar motion with Z axis support, a linear 2 axis system was created using guiding rods and bearings. By taking weight into account, the system used 6.35 mm precision ground 6061 aluminum rods and stainless steel linear ball bearings with aluminum housings (5911K11, NB corporations, Hanover Park, Illinois). For a 300-gr rat (1 Kg of total moving load) the max deflection of the guiding rod was calculated to be 3.023mm. The static and dynamic forces required to move the 2-axis system is reduced further by removing the linear bearings seals. Using the formula (equation 1) provided by the manufacturer (NB Corporation), the dynamic friction force of 0.0294 N/bearing was calculated after removing the seals.

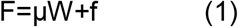

Where, F represents dynamic friction force (N), μ is the dynamic friction coefficient, W is the applied load (N) and *f* is the resistance from seal (N). Theoretical friction value of the linear bearings after removing the seal was calculated and then compared with the friction value experienced by the bearing during dynamic trials (Sup. table 2). To further define the static and dynamic friction forces of the BWS system, we conducted tests using an actuator (MT1-28, Thorlabs, N, New Jersey) and a load cell (Nano17, ATI, Apex, NC) (Fig. 1e,f). The actuator utilized micro-stepping and acceleration controls to provide an applied force to acquire friction and inertial data through the load cell across X and Y axis, and 45° angles.

#### Weight manipulation

As shown in Fig. 1c,d, the rodent’s weight can be manipulated by changing the Z axis position, variation of the pitch angle, and by shifting the rat’s weight bilaterally. The Z axis support uses a stepper motor and ball screw to accurately change the amount of body weight support provided in small increments.

### Surgical procedures

#### EMG wire and electrode implants for healthy and SCI rats

Nine (healthy: 6 and SCI: 3) adult female rats (Sprague–Dawley, 270–300 g body weight) were used in this study. The rats were deeply anesthetized by a combination of ketamine (100 mg/kg) and Xylazine (10 mg/kg) administered intraperitoneally (IP) and maintained at a surgical level with supplemental doses of ketamine as needed. Buprenorphine was administered as a single dose at the beginning of the experiment. All surgeries were performed under aseptic conditions. The experimental procedures complied with the guidelines of National Institutes of Health Guide for the Care and Use of Laboratory Animals and are conducted in accordance with protocol approved by the Animal Care Committee at the Mayo Clinic, Rochester MN.

A small skin incision was made at the midline of the skull. The muscles and fascia were retracted laterally, and the skull was thoroughly dried. A 12-pin Omnetics circular connector (Omnetics, Minneapolis, MN) and 12 Teflon-coated stainless-steel wires (AS632, Cooner Wire, CA) were attached to the skull with screws and dental cement as previously described^28,40^. Skin and fascia incisions were made to expose the bellies of the medial gastrocnemius (MG), and tibialis anterior (TA) muscles bilaterally. Using hemostats, the EMG wires were routed subcutaneously from the back incision to the appropriate locations in the hind-limb. Bipolar intramuscular EMG electrodes were inserted into the muscles as described previously^40^. The EMG wires were coiled near each implant site to provide stress relief. Electrical stimulation through the head-plug was used to visually verify the proper placement of the electrodes in each muscle.

A partial laminectomy was then performed at the L2 vertebral level (S1 spinal cord level) and one wire was affixed to the dura at the midline using 9.0 sutures as previously described ^28^ A small notch made in the Teflon coating (about 0.5–1.0 mm) of the wire used for EES was placed toward the spinal cord and served as the stimulating electrode. The wire was coiled in the back region to provide stress relief. Teflon coating (about 1 cm) was stripped from the distal centimeter of one wire that was inserted subcutaneously in the back region and served as a common ground^25^.

#### Spinal cord transection

One week after electrode implantation surgery, a complete spinal cord transection was performed on the three SCI rats. Rats were anaesthetized with a mixture of oxygen and Isoflurane (≈1.5%). Mid-dorsal skin incision was made between T6 and L4 and the paravertebral muscles were retracted as needed. A partial laminectomy was performed at the T8 level and the dura was opened longitudinally. Lidocaine was applied locally and the spinal cord was completely transected using a micro-scissors. Completeness of the lesion was verified by two surgeons under microscope. If so, tissues were sutured by layers and animals were allowed to recover in individual cages with soft bedding. Manual bladder expression was performed four times daily during two weeks post transection.

#### 6-OHDA injection

Three male Sprague-Dawley rats were intraperitoneally anesthetized with a mixture of ketamine (90mg/kg) and Xylazine (10mg/Kg) and received atropine methyl bromide (0.03mg/Kg, IM) and buprenorphine SR to reduce bronchial secretion and to provide 72 hours of pain relief, respectively. They were also pre-treated with Pargyline (50mg/Kg, IP) to prevent 6-OHDA catabolism by monoamine oxidase. Following induction of anesthesia, each rat was placed in a standard stereotaxic frame in which the skull was secured with a nose clamp, incisor bar, and ear bars. Body temperature was maintained constant at 37 degrees C by a heating pad. A single longitudinal incision of the scalp (2 cm in length) was made to expose the surface of the skull. A small piece of overlying cranium and dura was removed and a 26 gauge needle was stereotactically implanted into the right median forebrain bundle. 12μg of 6-OHDA solution (4μg/1μl) was infused at an injection rate of 0.25μl/min. The needle was left in place for additional 5 min after infusion to allow dispersion of the solution in the tissue and then was slowly removed. The wound was closed by suturing the subcutaneous fat and skin with absorbable 4.0 vicryl sutures using simple interrupted stitches.

### Data acquisition

Through fast synchronized communication using transistor-transistor logic (TTL), the motion analysis system, open field tracking camera, force transducer, electrophysiology acquisition unit and the stimulator can communicate and operate synchronously to collect multifactorial data and provide electrical stimulation of various parameters. For the SCI rats, EMG, kinematic, kinetic and open field video data were collected simultaneously once per week following 4 to 5 weeks post transection surgery. In control rats, data collection started 1 week post EMG wire implant surgery and in PD rats gait information has been collected 3 weeks after 6-OHDA injection. Custom MATLAB scripts were used to analyze the data.

#### Spinal Cord Electrical Epidural Stimulation

A single channel manually controllable isolated (A-M systems, Sequim, WA) or an eight-channel real-time programmable (STG4008, Multichannel Systems, Reutlingen) stimulators was used to deliver biphasic square wave pulses (250 μs pulse width) at 40 Hz with amplitudes ranging from 0.5 to 2 volts to the epidural electrode placed on the rat lumbosacral (S1) spinal cord.

#### Gait analysis

The motion tracking system (Vicon, UK) was used to record three-dimensional digital position of back and the hind limb joints (100 Hz). Six motion-sensitive Infra-Red (IR) cameras were aimed at the treadmill or the open field volumes. Another high-speed video camera synchronized with motion tracking system was positioned in a side to provide a lateral view of the motor performance. Retro-reflective markers were placed on bony landmarks at the iliac crest, greater trochanter, lateral condyle of the femur, lateral malleolus and the distal end of the fifth metatarsal on both legs of the rat to record the kinematics of the hip, knee and ankle joints. Nexus system was used to obtain three-dimensional coordinates of the markers. Gait cycles were defined as the time interval between two successive paw contacts of one limb. Successive paw contacts were visually defined by the investigators based on video records. Cycle duration, and stance and swing durations were also manually determined from the kinematic recordings. Each step of a sequence was resampled to average cycle duration of the steps collected during the session, thus minimizing elimination of temporal information. Limb endpoint trajectories were determined from motion of the toe marker. The computed step parameters included stance-swing phase, step frequency, stride length, maximum toe height and joint angles.

#### EMG and fMEP analysis

EMG activity was collected from TA and MG muscles at 4000 Hz during stepping and later high-pass filtered at 0.5 Hz to remove the DC offset. Synchronization pulse sent from motion tracking system was used to identify the same time window used in gait analysis. To perform fMEP (functional Motor Evoked Potential) analysis of EMG activity recorded during stepping, EMG signals were processed based on EES inter-pulse time windows, e.g. 25ms for 40 Hz and 10ms for 100 Hz stimulation, and organized sequentially. To quantify the number of mono and poly synaptic responses during 40 Hz stimulation, the numbers of peaks were counted within 5.5 to 9.1ms and 9.1 to 25ms. In order to determine flexor (TA) and extensor (MG) coordination, EMG signals were band pass filtered (10Hz to 1000Hz), rectified, normalized and plotted.

#### Kinetic recordings

A transducer (Nano17, ATI, Apex, NC) was used to record 6 measurements including force and torque across the X (*F_x_*, *τ_x_*), *Y* (*F_y_, T_y_*) and *Z* (*F_z_, T_z_*) axis. This load cell was placed directly above the rat’s centerline and measures force and torque produced by the rodent during locomotion tasks. A custom LabVIEW (National Instrument, Austin, Texas) script was used to acquire force and torque data. The signals were recorded at 100 Hz sampling rate and later high-pass filtered at 0.5 Hz to remove DC offset. Beginning and end of stepping phase was determined using the video recorded simultaneously. Average force and torque were determined dividing the total amount force with duration of recordings. Additionally, force vectors for each time points were determined using, F= *F_x_* i + *F_y_* j + *F_z_* k.

#### Open field locomotor tracking

To track movement and orientation of the rodent in an open field, video data were captured during trials. The open field video data were later analyzed using ImageJ (NIH) and custom MATLAB script (Mathworks Inc., Massachusetts) to obtain the two dimensional coordinates of rat’s position within open field. A custom MATLAB script was used to determine the distance travelled, average velocity and active time for each rats.

### Statistical analysis

Statistical analysis was performed using SigmaPlot (Systat Software Inc., UK). The data was first tested for normality and then one-way Analysis of Variance (ANOVA) was performed to determine if the groups had different outcomes. If the groups were found significantly different (p<0.05) a post-hoc analysis was performed using Holm-Sidak method to compare against control and p values were reported. When compared between two groups two samples t-test was performed. All results were reported as mean± standard deviation, and *p<0.05, **p<0.01 and ***p<0.001. To reduce the number of variables (n=34) measured using multiple modalities we performed multi-step statistical procedure Principal Component Analsysis (PCA) was performed on all the steps collected from different combinations of EES and pharmacological combinations (Sup. table 3). The first three PCs explaining the most amount of variance were used to plot the steps collected from all the configurations and visually demonstrate their clusters.

## Results

### Assessment of BWS system’s mechanical properties

After building the system, we calculated the theoretical dynamic friction values to the direction parallel to, Y axis (*F_y_*), diagonal to X and Y axis (*F*_45_), and across X axis (*F_x_*) for a 1 Kg load. The values of *F_y_*, *F*_45_ and *F_x_* were calculated to be 0.29 N, 0.12 N and 0.18 N, respectively. After building the BWS system, the experimental static and dynamic friction profile were obtained, while a constant force was applied (Fig. 1d-e). The maximum static and dynamic friction forces were observed across Y-axis (static: 0.28±0.1 N and dynamic: 0.14±0.05 N) and the minimum static and dynamic friction forces were found across X-axis (static: 0.16±0.02 N and dynamic: 0.09±0.03 N).

**Figure 1:**
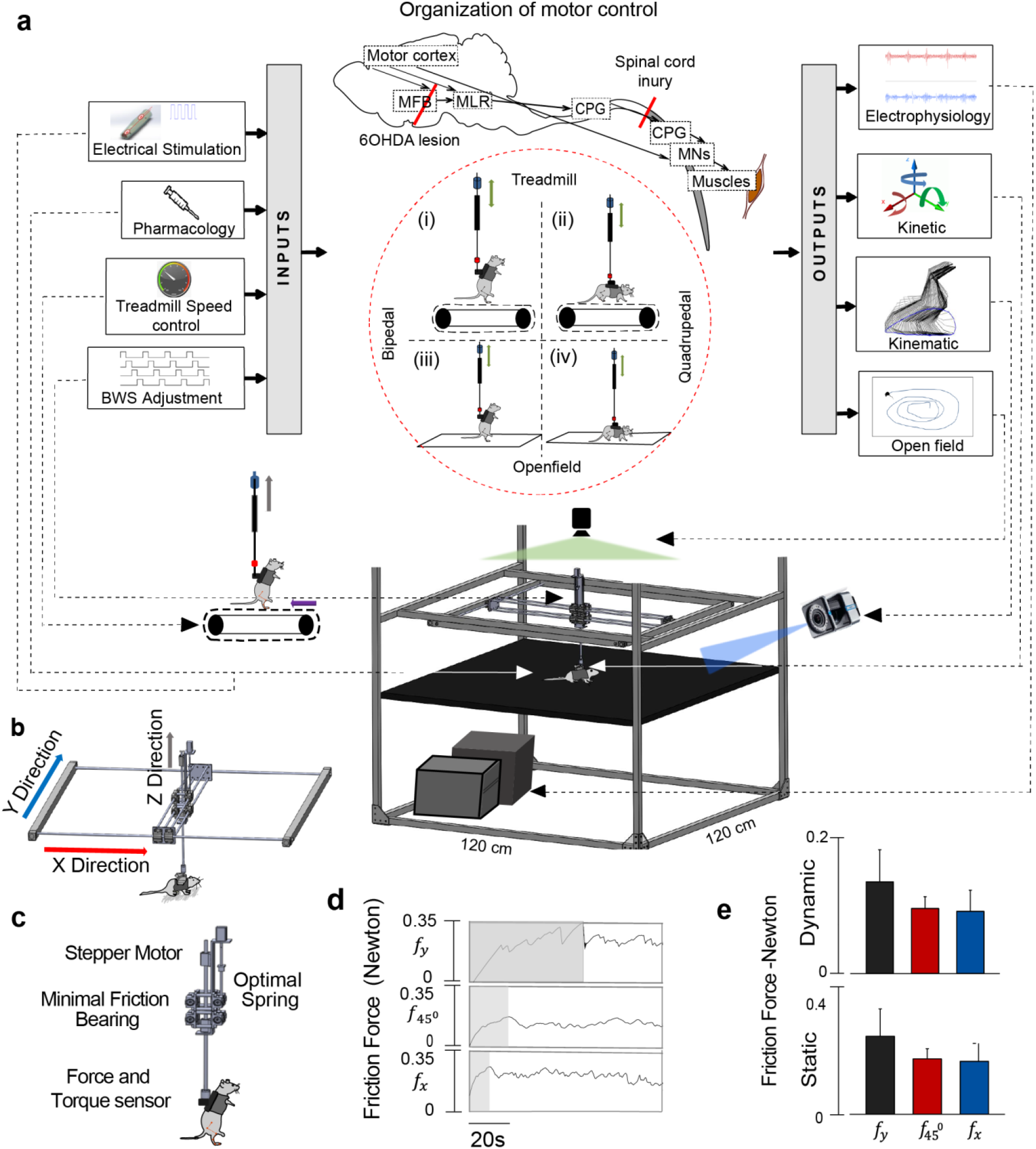
Diagram of the multifactorial behavioral assessment (MfBA) approach. Main inputs to the brain and spinal cord neural circuitry and related techniques to manipulate the neural networks though the changing of the afferent input, e.g. adjustment in electrical stimulation and pharmacological modulation, treadmill speed, and in BWS, presented on the left side of diagram. The main outputs related to the pattern of motor response generated by neural circuitry and recorded in the form of electrophysiological response (e.g. EMG, fMEP, etc.), kinetics, kinematic, and open field behavior presented on the right side of diagram (a). Brain and spinal cord main functional structures and related to this study brain and spinal cord lesions (red lines) are outlined on diagram between inputs and outputs. Main experimental conditions for assessment of motor behavior in this study included bipedal and quadrupedal stepping performed on a treadmill and in the open field also presented between inputs and outputs. Main components of the multimodal system include: BWS system, motion tracking system, open field camera, electrophysiology unit, stimulator, motorized treadmill and force, and torque transducer. (b) X and Y axis frames and central moving part of the BWS (c) Zoomed view of the central moving unit of the BWS showing components responsible for Z axis support and adjustment. (d) Friction force characteristic curves of the system. A 1.5 μN constant force was applied at X, 45°, and Y direction to obtain the friction characteristics. The shaded and non-shaded regions indicate the static and dynamic friction forces respectively. (e) Mean static and dynamic friction forces of the system (n=4).

### Validation of BWS system during treadmill and open field locomotion

Next, extensive analysis of gait parameters and open field behavior was performed while supported by the BWS in order to demonstrate the system’s performance during treadmill and open field locomotion.

#### (a) Treadmill stepping of neurologically intact and PD (6-OHDA) rodents

Gait parameters during treadmill stepping (treadmill speed: 15 cm/s) in healthy rats (n=5) while freely moving on a treadmill or supported in the BWS system during quadrupedal (~ 20% BWS) and bipedal (~60% BWS) locomotion were compared (Fig. 2a-c). Quadrupedal stepping after unilateral 6-OHDA injection in hemi-Parkinsonian rats (n=3, ~20% BWS, Fig 2d) were recorded and compared with the healthy rats. While freely running on a treadmill, healthy rats took 135.5±31.29 steps/min, but when supported by the BWS, they took 104.56±21.59 steps/min during bipedal stepping and 104.64±23.64 steps/min during quadrupedal stepping (Fig. 2e, free vs BWS-bipedal: p<0.001 and free vs BWS-quadrupedal: p<0.001). In contrast, PD rats took only 57.95±44.94 steps/min (BWS-quadrupedal vs. 6-OHDA PD Rat: p<0.001). At the same time, no significant difference was detected between stride lengths of free, BWS-bipedal, and BWS-quadrupedal rats (Fig. 2e, free: 19.03±6.23 cm, BWS-bipedal 16.91±3.6 cm, and BWS-quadrupedal 18.9±7.1 cm), although, BWS-quadrupedal stepping PD rats had smaller step lengths (14.4±4.8 cm) compared to BWS-quadrupedal healthy rats (BWS-quadrupedal 18.9±7.1 cm: p <0.001). Similarly, there were no significant differences between maximum toe heights during free, BWS-bipedal, and BWS-quadrupedal stepping (Fig. 2 g, free: 4.15±0.16 cm, BWS-bipedal 5.06±0.16 cm, and BWS-quadrupedal 4.84±0.16 cm). However, PD rats had significantly shorter toe height comparing to BWS-quadrupedal rats (6-OHDA PD rat: 2.25±.044 cm, p<0.001). In healthy rats, the stance phase was the shortest during freely stepping (56.68±7.73 %) and the longest during BWS-bipedal stepping (70.17±9.98 %, Fig. 2h). Both, BWS-bipedal and BWS-quadrupedal stepping had significantly longer stance phase (Free vs. BWS-bipedal: p<0.001 and free vs. BWS-quadrupedal: p<0.05). Additionally, 6-OHDA PD rats had significantly longer (p<0.05) stance phase compare to BWS-quadrupedal rats.

**Figure 2:**
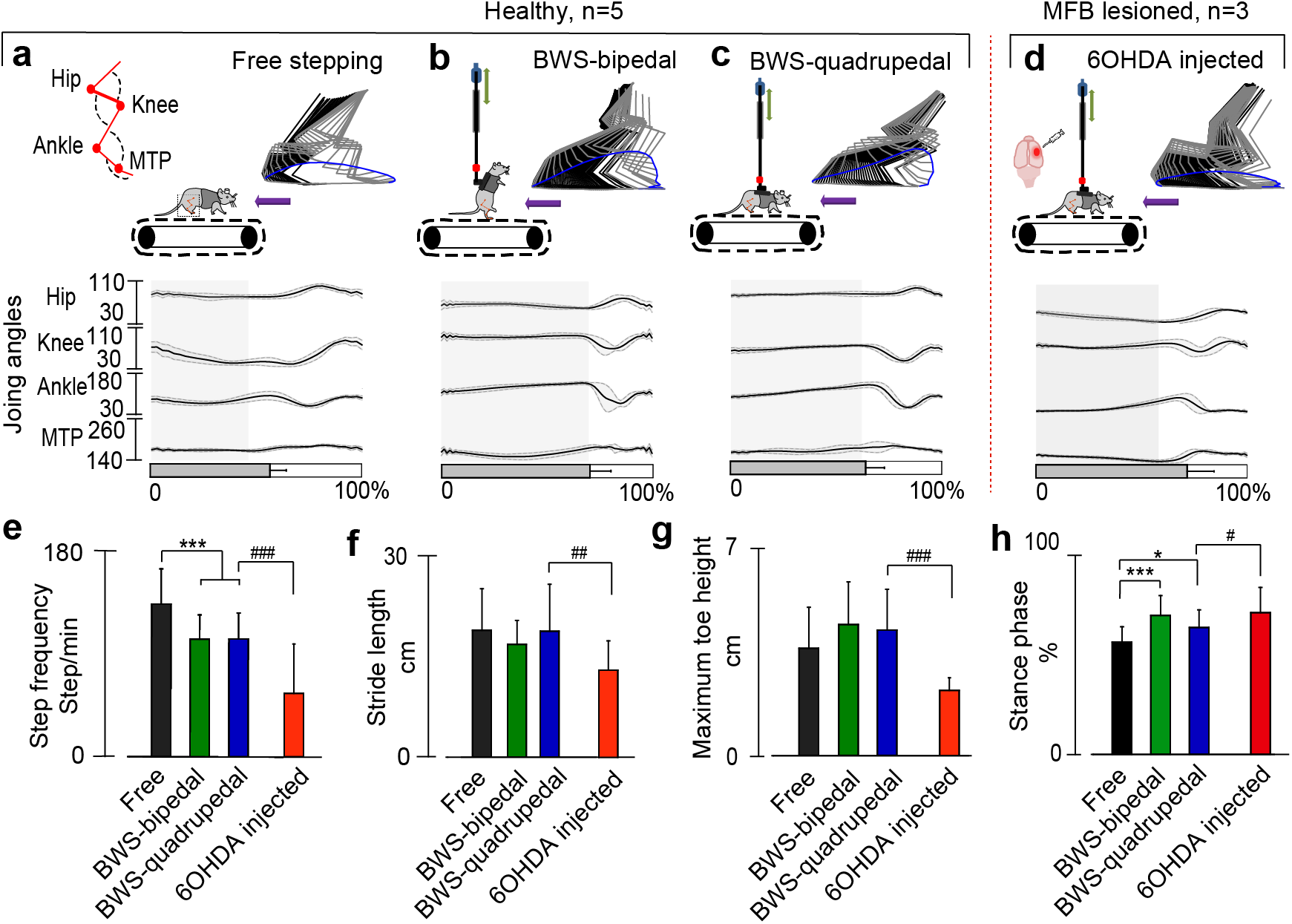
Kinematic assessment during bipedal and quadrupedal treadmill locomotion in healthy and PD rats. (a) Stick diagram, toe trajectory and joint angles (hip, knee, ankle and metatarsophalangeal (MTP)) of healthy rat running freely on treadmill (15 cm/s) for 10 consecutive steps. The shaded region represents the stance phase. The same rat was evaluated during (b) bipedal locomotion and (c) quadrupedal locomotion, while supported by BWS. A different rat with PD was evaluated 3 weeks after 6-OHDA injection also supported by the BWS (d). In all conditions involving BWS up to 20% of the rat’s body weight was supported. Comparison of gait parameters, (e) step frequency, (f) stride length, (g) maximum toe height, and (h) stance-swing cycle. (*p <0.05, **p<0.01 and ***p<0.001, one-way ANOVA, data represents mean± standard deviation, n=5 (healthy rats), n=3 (PD rats)).

#### (b) Simultaneous kinematic and open field behavioral assessment in neurologically intact rodents

To determine if the friction level of the BWS system is within tolerable range to allow open field locomotion in rodent, we performed open field test using healthy rats in the 80×80 cm area within the BWS system. The same rats (n=4) moved freely around the open field and also while supported by the BWS system for 10 minutes (Fig. 3a-d). The BWS system was found to significantly reduce the total distance travelled by the rats (Fig. 3b, freely moving: 29.72±4.03 m and BWS: 17.83± 9.16 m: p<0.05). However, the average velocity of the rats freely moving and moving in the BWS were not different (Fig. 3c, freely moving: 4.95±0.67 ms^-1^ and BWS: 3.24±1.51 ms^-1^), indicating limited dynamic friction force level. In addition, measuring the time rats were active in the open field (Fig. 3d) and found that freely moving rats spent 60.51±17.13 % of the time exploring the open field, while rats supported by the BWS spent less time, 42.9±8.54 %, although, this difference was not significant. Next, we compared freely-walking hind limb gait parameters to gait parameters collected during BWS quadrupedal open field locomotion in neurologically intact rodents (Fig. 3e-h). In order to allow quadrupedal locomotion, a maximum of 20% BWS was provided by the BWS system. Fig. 3e shows the representative gait of freely walking rats in the open field and also while supported by the BWS system. Hind limb stance and swing phases were not significantly different between freely walking rats (stance: 53.52±11.36 %) and BWS supported rats (stance: 52.29±8. 56%). Although rats in the BWS took more steps (173.22±36 steps/min) compared to freely walking rats (149.06±30.37 steps/min), that difference was not significant (Fig. 3f). Additionally, stride lengths (BWS: 4.21±2.26 cm and freely walking: 4.14±1.77 cm) were found to be similar (Fig. 3g). However, maximum toe height during BWS stepping (3.52±0.61 cm) was found to be significantly higher compared to freely walking (6.25±2.8 cm) (p<0.001) (Fig. 3h).

#### (c) Simultaneous kinematic and electrophysiological assessment during open field behavior

The feasibility of combining kinematic and EMG recording during open field locomotion was evaluated next (Fig. 3i). Open field trajectories were recorded continuously using a camera positioned above the open field platform, which was synchronized to limb kinematics recorded during two epochs (i and ii) (Fig. 3i). Using the open field trajectory, we quantified the maximum and average velocities of the rat during the two epochs of kinematic recordings (Fig. 3j) and found a maximum velocity (14.12 cm/s) during epoch-i, which was corroborated by a maximum step frequency (211.64± 17.71 steps/min) recorded by the kinematic system (Fig. 3l). We also quantified the peak-to-peak voltage of EMG traces recorded from the TA muscle (Fig. 3k) and found minimum peak-to-peak amplitudes occurred during epoch-i (2.91±0.3 mV). No significant differences were observed in strident length (Fig. 3m) and maximum toe height (Fig. 3n). However, significantly higher toe fluctuation was observed during epoch-i (9.83±3.86 cm compared to epoch-ii (4.03±1.4 cm) (Fig. 3o).

**Figure 3:**
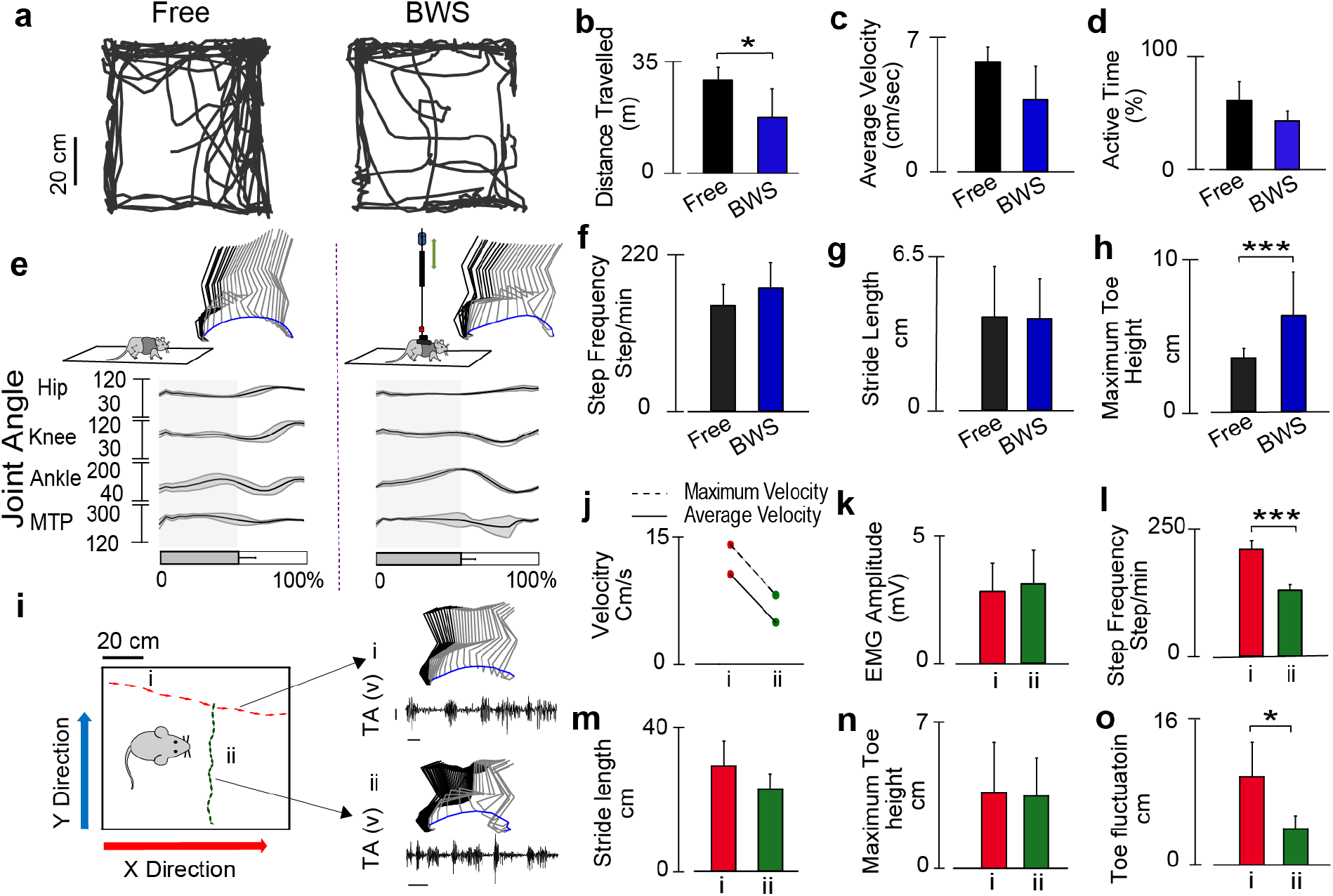
Evaluation of open field behavior and locomotion in the BWS system in neurologically intact rodents. (a) Open field trajectory of a freely moving healthy rat and the same rat supported by the BWS (~20%) during a 10 minutes evaluation. (b) Average total distance travelled in the open field, freely moving rat (black) and supported by the BWS (blue). (c) Average velocity (cm/sec) for the same two conditions. (d) Average of total active time by the rats (%). (e) Kinematic assessment during over-ground locomotion while freely walking and supported by BWS. Stick diagram, toe trajectory and joint angles (hip, knee, ankle and MTP) of healthy rat during locomotion on a runway for 8 steps collected during the same session. The shaded region represents the stance phase. Kinematics model is extracted using the ilium, hip, knee, ankle, MTP and toe markers positioned on the skin. Comparisons of gait parameters, (f) step frequency, (g) stride length and (h) maximum toe height. (*p <0.05, one-way ANOVA for n=4, data represents mean± standard deviation). (i) Example of open field trajectories, and simultaneously collected kinematic (stick diagram) and EMG traces (TA). A healthy rat was allowed to walk freely in the open field area while supported by the BWS and two representative epochs (i) and (ii) are shown here, which reflecting rat’s movement toward y and x directions, described in Fig. 1c. Scale represents 100ms and 1mV. (j) Maximum and average instantaneous velocity measured by the open field camera for the two epochs. (k) Peak to peak EMG amplitude of raw EMG (TA muscle), (l) Step frequency, (m) stride length, (n) maximum toe height and (o) toe fluctuation, of the same two epochs. (Bar plot represents mean± standard deviation).

### Variation in directional force and torque during treadmill locomotion in intact and SCI rodents

Ichiyama et al.^33^ and multiple following research reports using electrophysiological and behavior assessments described that, 0.3mg/Kg Quipazine (a 5HT2A/C agonist) injection can improve stepping outcome in SCI rat with or without EES. In this study we tested kinetic outcome of healthy and SCI rodent behavior with and without Quipazine. Using a six-axis force and torque sensor placed directly above the rat (Fig. 4a), we measured directional force and torque values produced during quadrupedal open field locomotion performed by a (i) healthy rat, (ii) same rat after complete SCI without any intervention, and (iii) after administration of Quipazine (Fig. 4b-d). Representative examples of directional force values, normalized force vector (*f_x_ i* + *f_y_j* + *f_z_ z*) and frequency spectrum analysis are presented in Fig. 4b-d. Healthy rats produced coordinated directional force values and force vector equally toward forward and reverse directions (marked by green and red arrows). After SCI the directional force values were less coordinated and mostly directed toward lateral and reverse direction, related to predominant dragging motion after SCI. Administration of Quipazine led in increase of force vectors toward forward and lateral directions. In healthy rat, frequency spectrum analysis demonstrated distinct frequency peaks in each direction, which mostly disappeared after SCI. After administration of Quipazine, both *f_y_* and *f_z_* peaks returned, although at lower frequency compared to healthy rat. The average force and torque produced during different trials (n=3) are presented on Fig. 4e. Forward force (*f_x_*) observed before injury was 0.35±0.25 N, where following SCI it decreased to 0.07±0.25 N. Interestingly, IP administration of Quipazine restored forward force to 0.49±0.2 N, that was higher compare to pre-injury level. At the same time, lateral force (*f_y_*) and vertical force (*f_z_*) were not different before and after SCI (f_y_, healthy: 0.33±0.37 N and SCI: 0.34±0.42 N, and *f_z_,* healthy: 0.95±0.68 N and SCI: 0.96±0.74 N). Both forces increased following administration of Quipazine (*f_y_*: 0.62±0.25 N and *f_z_*: 1.17±0.22 N). Similar to forward force (*f_x_*), left and right torque (*τ_x_*) decreased after SCI (*τ_x_*, healthy: 32.62±45.05 N-m and SCI: 22.23±22.28 N-m) and increased to the level higher than in healthy animal following Quipazine administration (53.59±15.32 N-m, Fig. 4e).

**Figure 4:**
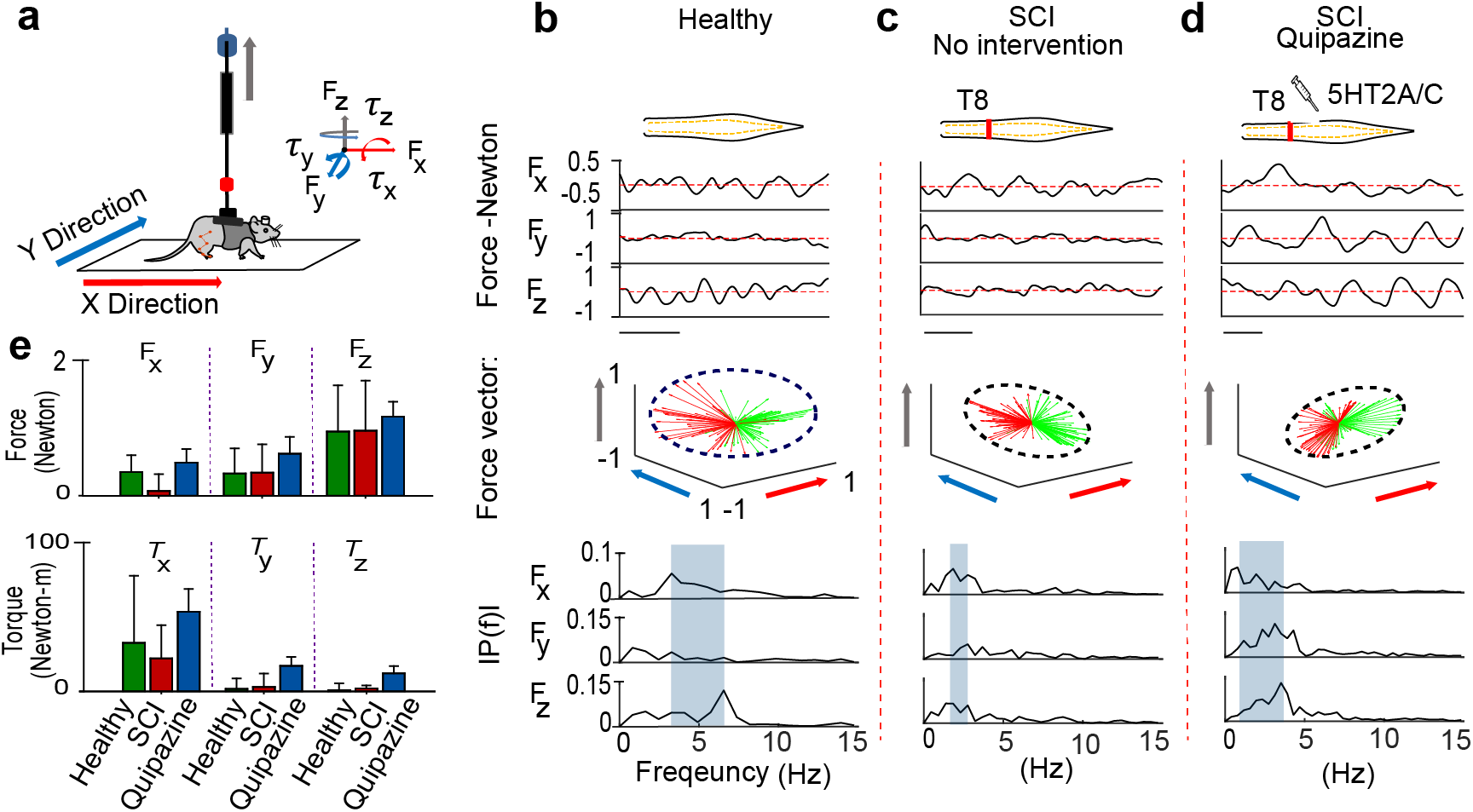
Directional force and torque during treadmill locomotion in intact and SCI rodents. (a) Experimental setup to collect kinetic information during rat’s locomotion in open field. Force and torque across X, Y and Z directions were collected. Representative example of force values, normalized force vectors and frequency domain response produced by, healthy rats (b), SCI rats without any intervention (c), and SCI rats after IP administration of Quipazine (0.3 mg/Kg) (d). The red dashed lines represent the average force (x axis scale: 0.5s). Force vectors with forward and backward directions are plotted with green and red arrows respectively. The outside edges of the force vectors were enclosed by the dotted ellipse in order to demonstrate their distribution. The axes of force vectors are indicated by the same color presented in Fig. 4a. The frequency ranges of peak force across three axes were demonstrated by the highlighted area. (e) Average force and torque values toward three directions recorded (n=3 trials). (Bar plot represents mean± standard deviation).

### Multi-factorial behavioral assessment of EES-enabled locomotion in rodents with complete SCI

Based on our previously published findings on modulation of spinal cord motor evoked potentials during standing^41^ and stepping^42^, we introduced evaluation of motor evoked potentials during task-related modulation of spinal circuitry (functional Motor Evoked Potentials or fMEP) in SCI rats 4 weeks after injury (Fig. 5a). Body weight supported multimodal system was designed for this study to provide a wide range of modulation of inputs and outputs to the spinal circuitry. Performed based on the number and amplitude of the selected components, fMEP analysis provides important information about modulation of the different components of spinal circuitry related to stepping.

To activate spinal locomotion circuitry, we choose 40 Hz epidural stimulation, commonly used in SCI rodents^23,27,28^. We applied two intensity level of stimulations, sub-threshold (80% of the motor threshold) and threshold EES that generates motor activity. Trajectory represents the rat’s whole body movement on a treadmill. The sub-threshold EES showed minimum to no movement, evident from open field and kinematic analysis. At the threshold level of stimulation, rat spent most of the time at the front of the treadmill. During sub-threshold stimulation, 40 Hz EES produced force vectors toward the right and left sides of the rodent; however, at motor threshold voltage intensities, force vectors shifted to forward and backward directionalities with respect to rodent’s body position.

EMG signals recorded from flexor (TA) and extensor (MG) muscles showed little to no alternating bursts during 40 Hz sub-threshold stimulation and alternating bursts, appeared during threshold level stimulation, along with increase in burst amplitude. Visualization of fMEP response of the step selected during EES-enabled stepping showed modulation of primary monosynaptic components during sub-threshold stimulation and only during period of TA activity, related to the swing phase of step cycle. At threshold level EES, monosynaptic components became larger and additional polysynaptic components appeared mainly in MG during stance phase of step cycle. Polysynaptic components in MG also followed a temporal pattern with gradual shifting (indicated by diagonal dashed line) as it was described earlier^42^, suggesting that modulation of the polysynaptic components of fMEP during EES is related to spinal activity responsible for coordinated activation of flexor and extensor muscles during stepping.

We quantified open field, kinematic, kinetic, EMG and fMEP variables (n=34 variables) across the modalities of output collected for both sub-threshold and threshold EES. As a large number of these parameters changed substantially between the conditions, in order to summarize their effect on output modalities, we performed principle component analysis (PCA). PCA identified the distribution of the steps facilitated by sub-threshold and threshold EES (Fig. 5b). The first three PCs explained 98% of the variance associated with the data. The sub-threshold and threshold steps found to be isolated in the PC space. In addition, sub-threshold EES steps were clustered together with minimum variation. In contrast, threshold EES steps were more spread across the PC space.

To demonstrate how the sub-threshold and threshold EES-enabled stepping vary across multiple modalities, we also presented direct comparison between tested modalities (Fig. 5c-g). This allowed to demonstrate how the higher level motor outputs (e.g. open field and kinematic parameters), correlates with functional state of spinal cord circuitry (assessed with fMEP) during tested behaviors. Increasing EES intensity from sub-threshold level to threshold level resulted in significant increase in distance travelled by the rat (Fig. 5c, p<0.001), improved forward force (Fig. 5d, P<0.01), and increased toe height (Fig. 5e, p<0.001). EMG amplitude in both TA and MG muscles also increased significantly (p<0.001), although these changes were more prominent in MG (Fig. 5f). In order to evaluate the contribution of polysynaptic components of spinal cord circuitry, during sub-threshold and threshold EES, we quantified the number of polysynaptic peaks during these two conditions (Fig. 5g). Polysynaptic peak counts did not change significantly during sub-threshold and threshold EES in TA, but increased significantly in MG at threshold level, indicating predominant influence of EES at threshold level on activity in extensor muscles during stance phase (p<0.001), reflecting circuitry modulation by sensory input during step cycle. This multifactorial analysis of simultaneously collected multimodal data shows a direct correlation between alteration of the states of spinal cord locomotor circuit and subsequent effect on locomotor behavior.

**Figure 5:**
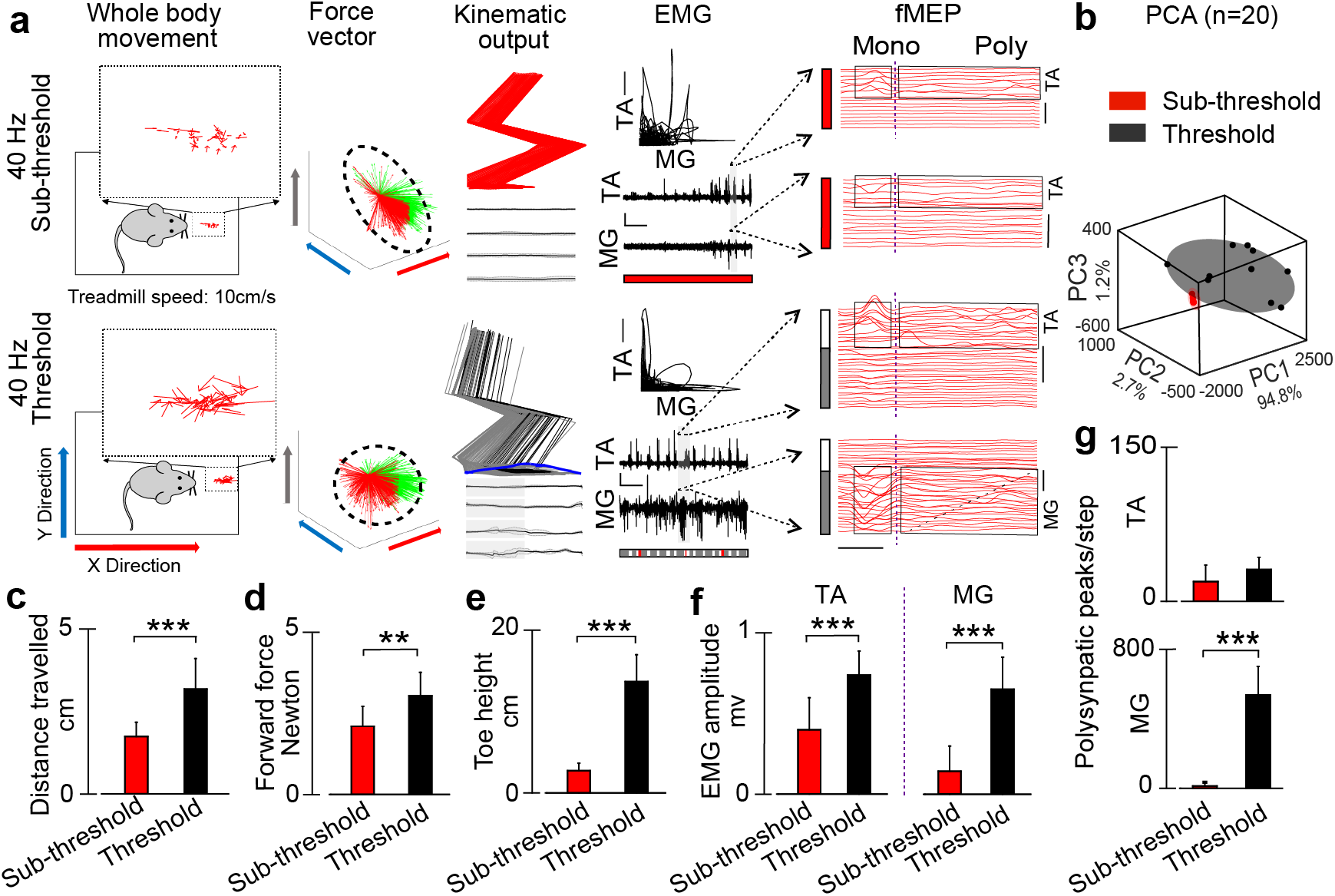
Multifactorial behavioral assessment in SCI rodents. (a) Bipedal stepping on treadmill (speed: 10 cm/s) induced by 40 Hz sub-threshold (~0.55 volt) and threshold (0.8mV) EES, in SCI rat 3 weeks after T8 transection. MfBA was performed based on several outputs: open field trajectory, force vectors, kinematic, EMG response with muscles antagonist’s correlation, and fMEP with analysis of individual motor evoked responses performed during stepping facilitated with different stimulation intensities. Grey area on EMG indicates the interval taken for fMEP analysis. Dash-line rectangle on fMEP highlights monosynaptic and polysynaptic components of fMEP. Scale: EMG, 1s and 2.5mV. fMEP, time: 5ms and amplitude: 0.5mV, and fMEP-AVG, amplitude: 0.1mV. On fMEP analysis, the lowest line corresponds to the earliest time point. (b) Principal component (PC) analysis was applied on the measured variables (n=34) across all tested modalities (for details see Sup. table 3). The steps (n=20) collected during subthreshold and threshold level stimulation are represented in the new three-dimensional (3D) space created by the three first PCs (explained variance, 98%). Average distance travelled (c), forward force (d), toe height (e), EMG amplitude (f), and fMEP peak counts in TA and MG muscles (g), during sub-threshold and threshold 40Hz stimulation presented. (*p<0.05, **p<0.01 and ***p<0.001, one-way ANOVA, n=10 steps in each condition. Data represents mean± standard deviation).

### Multifactorial behavioral assessment of EES and pharmacologically-enabled locomotion in rodents with complete SCI

The MfBA of spinal neural circuitry responsible for locomotion based on behavior and electrophysiological features was further extended in SCI rats (n=3) 4-5 weeks after complete T8 transection under a combination of EES and pharmacology facilitated stepping (Fig. 6). Although, specific kinematic characteristics of these synergistic neuromodulators have been discussed, previous studies^27^ failed to correlate the behavioral outputs, such as changes in kinematic, kinetic, and open field, with the state of spinal cord locomotor circuit reflected by electrophysiological markers, like mono- and polysynaptic evoked responses collected concurrently. Our goal was to further characterize this relation, thereby investigating the mechanism of the effect of spinal cord neurostimulation. The following conditions were compared: sub-threshold level of EES (40 Hz) in healthy rat, threshold level of EES (40 Hz) in SCI rats, sub-threshold level of EES (40 Hz) with IP administration of Quipazine in SCI rats, and threshold level of EES (40 Hz) with IP administration of Quipazine (Fig. 6b-f) in SCI rats. Sub-threshold level (<0.6 V) of stimulation was applied in healthy rats in order to obtain fMEP without influencing motor output during bipedal stepping on the treadmill while supported by the BWS (60% BWS). Healthy rats did not show any sign of discomfort during sub-threshold stimulation and exhibited regular pattern of stepping (Fig. 6b). Following SCI, no stepping was observed during sub-threshold EES (Fig. 6c). EES at threshold level alone (Fig. 6d), at subthreshold level with Quipazine injection (Fig. 6e), and at threshold level with Quipazine injection (Fig. 6f), produced consistent stereotypical steps. The number of steps taken per minute decreased significantly from healthy rats (76±14 steps/min) to SCI (0 steps) rats when subthreshold EES was applied in both groups (Fig. 6g, p<0.01). Threshold EES (82.63±19 steps/min), sub-threshold EES with Quipazine (62.18±10.57 steps/min) or threshold EES with Quipazine (62.42±8.81 steps/min), significantly increased step frequency and restored it to healthy level (p<0.01). Similar to step frequency, maximum toe height decreased from 7.49±2.7 cm to 0.13±0.04 cm after SCI when sub-threshold EES was applied (Fig. 6h, p<0.001). Threshold EES significantly increased maximum toe height (24.17±8 cm, p<0.001); however, highest toe height in SCI rats were observed during sub-threshold (51.43±8 cm) and threshold EES (53.64±8.7 cm) with Quipazine (p<0.05).

EMG activity recorded from TA and MG muscles of SCI rats was evaluated under the same conditions (Fig. 6i). During sub-threshold EES, SCI rats produced minimal to no coordination between TA and MG muscles. However, at threshold level EES, coordination between TA and MG muscles was evident. Quantification of peak to peak EMG amplitude in both TA and MG muscles showed, significant reduction following SCI when sub-threshold EES was applied (p<0.001) (Fig 6j).Threshold EES, sub-threshold EES with Quipazine or threshold EES with Quipazine significantly increased TA amplitude (p<0.01). Interestingly, in MG muscle, even though threshold EES restored amplitude to preinjury level (p<0.001), sub-threshold EES with Quipazine injection failed to increase amplitude. However, threshold EES with Quipazine injection was effective in increasing MG amplitude (p<0.001).

Fig. 6k provides example of modulation of mono- and polysynaptic components of fMEP recorded from TA and MG muscles for the steps highlighted with grey area in the EMG burst presented in Fig 6i. In healthy and SCI rats, Mono- and polysynaptic activity showed distinct spatial and temporal distribution, when modulated with sub-threshold and threshold EES, with or without Quipazine injection. In order to identify how EES with and without Quipazine modulates spinal cord polysynaptic activity, reflecting modulation of spinal circuitry generating characteristics motor output, we quantified the number of polysynaptic peaks and the area under curve (AOC) for both TA and MG muscles (Fig. 6l-m). In TA muscle, healthy rats with sub-threshold EES generated ~400 polysynaptic peaks/step, which decreased to ~12 peaks/step after SCI when same sub-threshold EES was applied (p<0.01, Fig. 6l). Threshold EES failed to significantly increase number of polysynaptic peaks. However, sub-threshold EES with Quipazine injection or threshold EES with Quipazine injection, significantly increased polysynaptic peaks (~1000 and ~700 peaks/step, correspondingly; p<0.001). Interestingly, subthreshold EES with Quipazine injection increased polysynaptic peaks/step even higher than healthy rats with sub-threshold EES (p<0.05). MG muscle of healthy rat during sub-threshold EES produced ~500 peaks/step, but following SCI the number of polysynaptic peaks decreased to ~10 peaks/step (p<0.001). Threshold EES increased the number of polysynaptic peaks to ~500 peaks/step (p<0.001). However, sub-threshold EES with Quipazine injection failed to increase the number of polysynaptic peaks/step to ~80 peaks/step; however, threshold EES with Quipazine injection did increased the number of polysynaptic peaks (1100 peaks/step, p<0.001), even higher than healthy rats with sub-threshold EES (p<0.05).

In TA muscle, AOC did not change after SCI with sub-threshold or threshold EES (Fig 6m). In SCI rats, Quipazine injection with threshold EES increased AOC higher than healthy rats with sub-threshold EES or SCI rats with sub-threshold or threshold EES (p<0.001). In MG muscle, AOC also did not change after SCI when sub-threshold EES was applied. Threshold EES however, significantly increased AOC (p<0.001), which decreased again when Quipazine was injected during sub-threshold EES (p<0.001). Threshold EES with Quipazine did increase MG AOC compared to sub-threshold EES with Quipazine (p<0.01), but this was still lower than threshold EES (p<0.001).

10 fMEP variables were quantified (Sup. table 3) from TA and MG muscles from the 100 steps collected under the same five conditions. In order to show, if these steps can be separated based on the variance demonstrated by the fMEP variables, we have performed PC analysis. The results of the PCA are presented in the 3D graph in Fig. 6n, by plotting the first three PCs. These PCs explain 85.3% of the variance associated with the fMEP variables. The highlighted elliptical spaces represent steps taken under different conditions. To clearly demonstrate the findings of PCA analysis, we have plotted the PCs in 2D views representing planes PC1 vs PC2, PC1 vs. PC3 and PC2 vs. PC3. The steps collected from different conditions maintain their clusters in the 3D and all three 2D views. The largest isolation between different groups was observed in the view perpendicular to the plane PC1 vs. PC2. Healthy rats with subthreshold EES (black ellipse) produced steps which are clustered together in 3D space and also on PC1 vs PC2 plane. These two PCs together explain 67% of the variance associated with the data. After SCI, steps produced with sub-threshold EES (red ellipse) move to the bottom of 3D space and form a small cluster. Threshold EES (yellow ellipse) and sub-threshold EES with Quipazine (pink ellipse) enabled steps to the opposite directions in PC1 vs PC2 plane and form two clusters. When threshold EES is applied with Quipazine the resulting steps (green ellipse) steps are produced in the middle of these clusters, indicating synergistic effect of threshold EES and Quipazine to facilitate stepping. PC analysis also showed that threshold EES enabled steps represents the closest to healthy rats electrophysiological pattern of fMEP compare to stepping facilitated with Quipazine applied at sub-threshold or at threshold level of EES.

**Figure 6:**
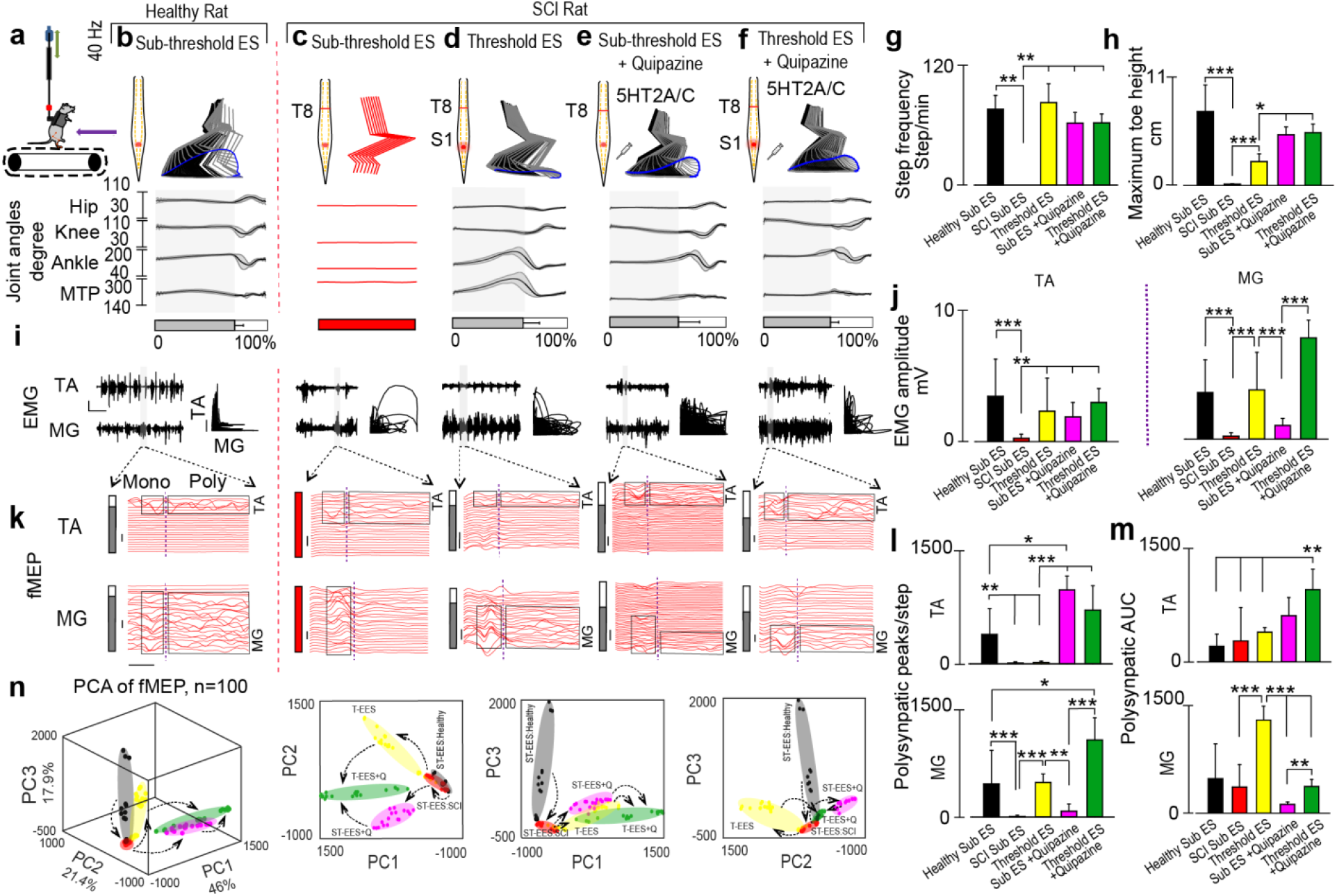
Multifactorial behavioral assessment of electrical and pharmacological modulation in SCI rodents. (a) Diagram showing the experimental settings with rat being supported by the BWS. The amount of BWS for SCI rats were kept constant at 85-90% for all conditions on the treadmill (speed=11.66 cm/s). (b) Representative treadmill stepping performed by healthy rats during sub-threshold 40Hz EES, (c) SCI rat with sub-threshold 40Hz EES, (d) SCI rat with threshold level 40Hz EES, (e) SCI rat with IP administration of Quipazine (5HT2A/C agonist) combined with sub-threshold 40Hz EES and (f) SCI rat with IP administration of Quipazine (5HT2A/C agonist) combined with threshold 40Hz EES. The quality of stepping is presented by stick diagram, toe-trajectory and joint angles (hip, knee, ankle and MTP). The shaded region indicates the stance phase. Average step frequency (g), and maximum toe height (h), for the same conditions are presented. (i) EMG bursts collected from TA and MG muscles during same configurations. Scale represents 1s and 2.5mV. Normalized coordination between TA and MG muscles presented for the same steps. Scale represents 0.25 Arbitrary Unit, where x and y axes extents from 0 to 1. (j) Peak to peak amplitude of TA and MG muscle for the experimental conditions. (k) fMEP of the step highlighted by grey area in the EMG signals in Fig. 5i for the conditions for both muscles. The area enclosed by black rectangles represents the mono and poly synaptic components of fMEP separated by the dashed line. fMEP scale: 5ms and 2.5mV. Number of poly synaptic peaks/step (l), and polysynaptic response’s area under curve (AUC) (m), in TA and MG muscles for the five conditions are presented. (n) Principal component (PC) analysis was applied on the derived fMEP variables (n=10) (for details see Sup. table 3 fMEP section). Total 100 steps were collected during the five conditions are represented in the new three-dimensional (3D) space created by the three first PCs (explained variance, 85.3%), and in the 2D planes. The steps under the same condition were highlighted by arbitrary elliptical shapes representing their clusters. The 2d views of the principle components are also presented to provide details of distribution of the steps collected under different configurations. Arrows indicate the consecutive changes in fMEP from healthy state to SCI (1), in SCI rodents from sub-threshold level of EES to threshold level of EES (2), and from sub-threshold level of EES to Quipazine administration (3), or to combination of EES and Quipazine (4) (*p<0.05, **p<0.01 and ***p<0.001, one-way ANOVA, n=3 SCI rats. Data represents mean± standard deviation).

## Discussion

Traditionally, rodent locomotor activity after SCI has been assessed using the Basso-Beattie-Bresnahan (BBB) open field locomotor rating scale, which summarizes the extent of motor recovery and has been correlated to lesion severity^43,44^. At the same time, the BBB and similar measures^45–47^ only account for a few basic movement characteristics while lacking sensitivity to locomotor variability and electrophysiological outcomes^48^. In response to these limitations, more sensitive methods to measure motor recovery, using kinematics, kinetics, and EMG^49,50^ were employed. Here, we introduced conceptually new approach to evaluate open field and treadmill-based locomotor activity in healthy and motor impaired rodents supported by a BWS simultaneously with evaluation of electrophysiological characteristics of spinal circuitry responsible for rhythmic motor output. Unique features of this approach and designed BWS system include integrated spinal stimulation, electrophysiological, and biomechanical recordings during *in vivo* motor activities. The integration of these experimental factors makes it feasible to correlate the *in vivo* activity of spinal circuitry with changes in motor behavior. Using MfBA approach, we found that characteristics of polysynaptic spinal activity in flexor and extensor muscles are critical in achieving coordinated stepping. MfBA that integrates experimental variables of this study particularly indicates on the importance of polysynaptic network activity during EES-enable locomotion that would not be evident with other approaches.

To validate designed for this study BWS system, we performed extensive mechanical tests to determine the friction forces imposed on rodents by BWS components of the system. The maximum dynamic frictional force of 0.28±0.1 N in the Y direction is within the range of propulsive force (~± 2N/Kg) produced by healthy rats during locomotion^51^. Additionally, the system’s static and dynamic friction values are in line with values of Rabinowicz’s sticking and sliding model, which describes the friction profiles of two lubricated sliding metal surfaces^52^. Therefore the system we developed imposes minimal extrinsic force during motor task performance in contrasts with recently developed, robotically-assisted BWS systems that are comprised of large, expensive structural and mechanical components and contain active serial actuators, which convolute results through inherent integration of intrinsic friction forces produced by the system and forces produced by SCI rodents during locomotion^53^. Some earlier versions of rodent BWS system relies on attaching robotic arms to the rat’s hindlimb in order to measure hindlimb trajectories^54,55^. However, this method is only suitable for less severe injury and may fail to adapt with the sudden change in locomotion.

During system validation, we characterized control bipedal, control quadrupedal, and PD quadrupedal treadmill stepping (Fig. 2). As expected, contralateral to lesion left hind limb gait deficits were identified in the PD rats. Specifically, PD rats exhibited reductions in step frequency, stride length, and maximum toe height, with increased stance phase duration (Fig. 2e-h). Previous studies also reported impairments in contralateral hind limb of unilateral 6-OHDA – rats in the form of shorter steps with reduced toe clearance^56,57^, which reflects the shuffling gait in human PD patients^58^. It has been reported that, pattern-generating networks responsible for stepping in dopamine-depleted rats are still functional and can produce coordinated rhythmic gait pattern^59^. A novel assessment system capable of providing time varying sensory input while simultaneously recording motor behavior can be further deployed to study the mechanism of motor deficit in rodents with brain lesion and for evaluation of therapeutic options.

In this study several functional tests, including open field assessment of neurologically intact rodents with BWS were used to validate system performance and MfBA approach. We found that rodents explored less in the BWS compared to freely exploring (Fig. 3b), however, average velocity and duration of activity were not significantly different between freely exploring rats and rats in BWS, indicating that BWS did not impede motor activity. We also characterized open field gait parameters of healthy rats (Fig. 3e-h). During over-ground stepping increased toe height was observed during BWS supported locomotion compared to freely exploring rats (Fig. 3h). Other gait parameters (e.g. step frequency, stride length) were not statistically different between freely walking BWS supported stepping, which also supports the conclusion that the BWS system did not hinder gait. The differences in toe height during BWS versus free open field locomotion were not observed during treadmill locomotion (Fig. 2g), indicating that kinematic parameters of open field locomotion may be more sensitive to perturbations, and in turn, may prove useful as targeted gait characteristics to enhance recovery via emerging therapeutics. In contrast, treadmill training is less prone to external perturbations yet provides the opportunity to investigate repetitive gait characteristics at investigator-controlled presets.

The system’s capability of recording locomotor kinetics using a force and torque transducer integrated into the BWS apparatus was demonstrated (Fig. 4, 5). Without any intervention, SCI rodents generated substantial forces in lateral directions, but only minimal forces forward and backward, which are essential in order to propel during locomotion. Following Quipazine (5-HT_2A_ and 5-HT_3_ receptors agonist) administration, forward and reverse directional forces increased to levels that were higher than those recorded prior to SCI (Fig. 4e). Lumbosacral EES produced stepping characterized by low toe height clearances during swing phase with increased step frequency while the administration of Quipazine without EES increased toe height and reduced stepping frequency. At the same time, Quipazine and EES synergistically modulated spinal cord networks to enhance treadmill stepping performance after SCI (Fig. 6). Previous studies have shown that EES enabled a greater number of steps compared to just Quipazine administration^33^. Additionally, previous reports found that Quipazine-induced stepping was dose-dependent with respect to achieving higher toe height during swing phase compared to Quipazine and EES combined stepping^33^, suggesting the synergistic effect could be due to an increased state of excitability within spinal sensorimotor networks achieved by Quipazine, which can be modulated by afferent feedback and EES to induce robust motor outputs and improved stepping performance^33^. Quipazine and other frequently used pharmacological agents in SCI animal models are not approved for use in humans and, unfortunately, cannot provide a clear impact on effect of neuromodulation in clinical settings at this moment. Additionally, the short time duration of their effects requires continuous administration and the long-term systemic effects of Quipazine remain unclear. Kinetic assessment in combination with open field tracking, EMG and fMEP assessment as presented in this study can be used to study the effect of EES alone or in combination with pharmacological facilitation. The BWS system designed in this study is capable of evaluating the effect of EES with additional pharmacological facilitation in various experimental settings. Since the combined effects of Quipazine and EES cannot be examined in humans, it will be critical to establish clinically-oriented assessment paradigms, similar to the MfBA developed for rodents, in order to achieve optimal EES-enabled motor performance in humans.

Established in this study characteristics of mono- and polysynaptic components in flexor and extensor muscles during EES synchronized functional tasks in healthy and SCI rats indicate on significant alteration of motor outputs (e.g. open field movement, kinematic and kinetic output). Associated with alteration of mono- and polysynaptic components recorded from related muscles during EES synchronized functional tasks, modulation of mono- and polysynaptic components in this study for the first time provided integrative approach to evaluate spinal locomotor circuitry and resultant motor behavior simultaneously. Specific changes in polysynaptic activity observed between tested groups could reflect complex changes in spinal circuitry. Thus, decreased number of polysynaptic components of fMEP after SCI was successfully compensated by administration of EES alone or with Quipazine, although EES alone was able to compensate polysynaptic activity only in MG muscle active during stance phase when circuitry related to MG receiving specific sensory facilitation from the foot, while component of circuitry related to TA could has limited sensory facilitation during swing phase and accordingly may require additional pharmacological modulation to became sensitive to EES. These findings suggest that fMEP integrated with MfBA approach could be a powerful tool in evaluation of spinal circuitry and related *in vivo* motor behavior.

In recent clinical studies^11–13^, a wide range of stimulation parameters were used to produce activity-dependent modulation of spinal sensorimotor networks, indicating a clear lack of information available to investigators during stimulation parameter optimization. Additionally, it remains unknown how various EES parameters influence human spinal circuit activity and longterm plasticity over the course of motor training. Currently available systems for motor behavior assessment, unfortunately, do not support necessary integration and real-time visualization of multiple assessments of spinal cord inputs and outputs^36–39,54^. In order to optimize EES-enabled activity-dependent modulation of human spinal network in an efficient and effective fashion, the faciliatory effects of EES on locomotor circuitry must be studied using integrative clinical assessment tools, which could be further design with MfBA approach similar to the system we have developed for rodent use.

In summary, by combining multiple assessment parameters including kinematic, kinetic, open field, and electrophysiology during therapeutically-enabled locomotion, we have established and tested a comprehensive real-time evaluation of motor behavior in healthy rodents and in rodents with neurologic deficits at the different CNS levels that significantly impact motor activity. Additionally, we provide evidence that MfBA combined with fMEP analysis is effective tool for at dissecting therapeutically-enabled gait characteristics and at targeting of different components of locomotor circuitry related with variations in motor behavior. Novel MfBA approach and designed for this purpose BWS system provide a platform for future investigations of the interactions between CNS inputs and outputs while manipulating external perturbations, and therapeutic administration in order to better understand how the CNS coordinates and executes complex motor tasks in healthy animals and in animals with neurologic deficit at different CNS levels.

**Supplementary table 1:** Components and cost of the BWS system.

| Part | Quantity | Price (\$) | Total (\$) |
| --- | --- | --- | --- |
| Corner cubes | 4 | 6.8 | 27.3 |
| 8020 Aluminum extrusion | 2 | 31.5 | 63.1 |
| Adjustable feet | 4 | 13.3 | 53.4 |
| Button head bolt | 12 | 11.6 | 11.6 |
| Steel rod | 2 | 15.7 | 31.5 |
| Aluminum Rod | 2 | 25.6 | 51.3 |
| Linear bearing | 4 | 13.9 | 55.8 |
| Stop collars | 9 | 3.8 | 35.0 |
| Rubber cushion | 8 | 1.6 | 13.5 |
| rod mount bolts | 8 | 0.1 | 1.0 |
| Spring | 1 | 8.7 | 8.7 |
| Miscellaneous | - | - | 40 |
| **Total price:** | - | - | **~ 400** |

**Supplementary table 2:**
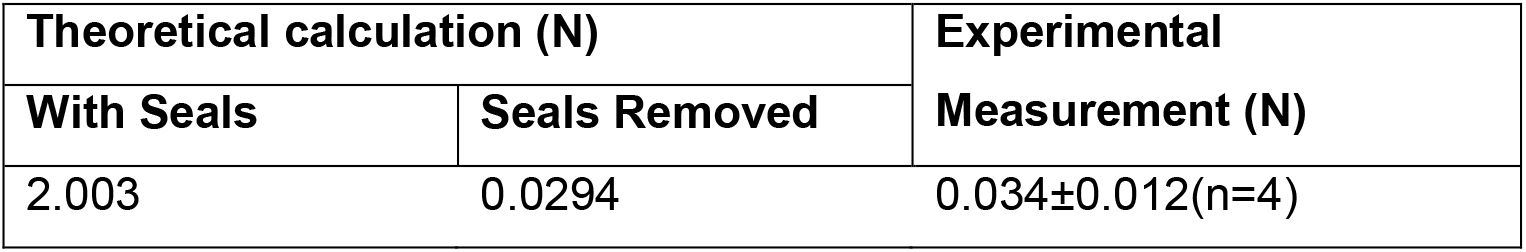
Theoretical and experimental frictional force analysis of a single linear

**Supplementary table 3:**
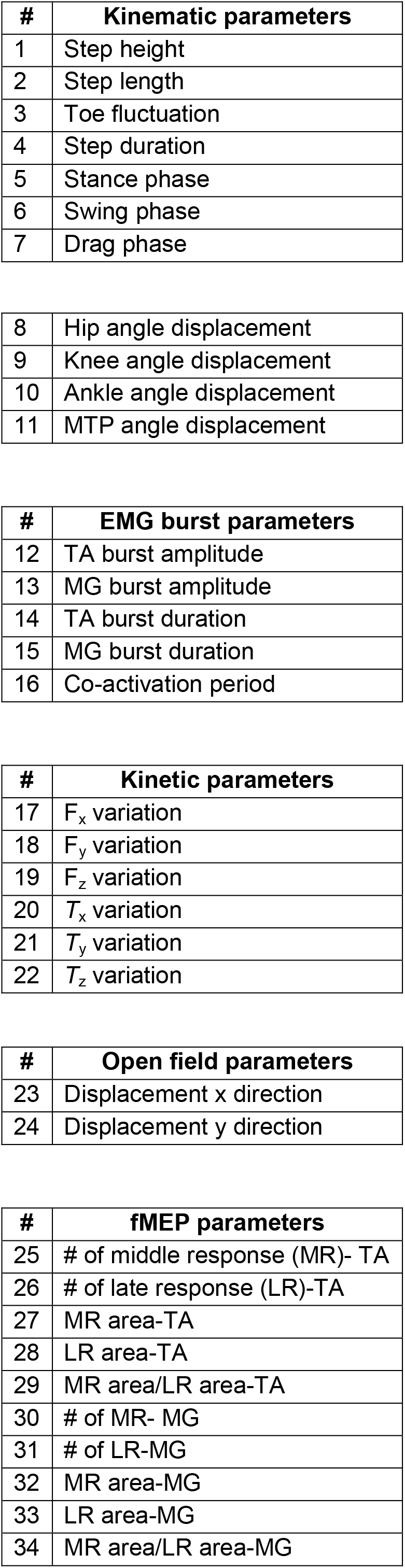
Summary of computed kinematics and EMG parameters for PCA.

## Acknowledgements

The authors thanks Mr. Seungleal (Brian) Paek for surgical and experimental assistance related to the Parkinsonian model. Finally, the authors thank Mayo Clinic’s Division of Engineering for their support to design and construct the multifactorial assessment system.

